# Machine Learning Analysis of Cilia-Driven Particle Transport Distinguishes Primary Ciliary Dyskinesia Cilia from Normal Cilia

**DOI:** 10.1101/2025.11.02.686130

**Authors:** Nicholas Hadas, Huihui Xu, Shambhawee Neupane, Wang K Twan, Ahmed Elgamal, Amjad Horani

## Abstract

**Rational:** Primary ciliary dyskinesia (PCD) is a genetic condition that results in dysmotile cilia and abnormal mucociliary clearance. Despite advances in understanding the pathogenesis of PCD, diagnosis continues to be challenging. Here we used feature-based machine learning and image-based deep learning to objectively quantify the directed particle transport of motile cilia and detect PCD-related cilia dysfunction.

**Methods:** Fluorescent microspheres were captured on cultured multiciliated cells using high-speed video microscopy as a proxy for motile cilia function. An interactive Jython script was designed to automatically detect, track and extract raw track metrics from videos. Data was subsequently analyzed to approximate a quantifiable and visual signature of ciliary transport through a custom-built Python Package, CiliaTracks.

**Results:** Airway epithelial cells were obtained from 14 individuals with genetically confirmed PCD, 10 healthy donors, and 2 patients with cystic fibrosis. A total of 602 videos (301 PCD and 301 non-PCD) were captured. Quantitative and visual analyses of fluorescent microsphere trajectories, including kinematic metrics and trajectory plots, revealed distinct motility profiles between PCD and non-PCD samples. Classical machine learning models and a convolutional neural network were employed to classify PCD using both modalities, demonstrating excellent accuracy of 95-97%, and the capacity to differentiate PCD from normal cells or cystic fibrosis.

**Conclusion:** Cilia-propelled microsphere transport exhibits unique trajectory patterns in PCD, enabling differentiation from non-PCD samples. Machine learning provides an objective and accurate framework for characterizing ciliary dysfunction, offering potential as a diagnostic tool for PCD.

## Introduction

Primary Ciliary Dyskinesia (PCD) is a rare genetic disorder caused from structural and functional abnormalities of motile cilia [1]. In the airways, motile cilia are critical for clearing pathogens and transporting fluids [2]. Ciliary dysfunction leads to a wide range of debilitating symptoms, including recurrent sinopulmonary infections, infertility, and organ laterality defect [3, 4]. The genetic basis of PCD is heterogeneous, with pathogenic variants in over 60 different genes identified as causative for PCD [5]. These disruptions typically result in cilia that are dyskinetic or immotile, which compromise the coordinated, propulsive movements essential for maintaining normal ciliary clearance. Normal ciliary transport is characterized by a high degree of coordination and directionality, as cilia beat in metachronal waves to propel mucus in a uniform path [6, 7]. The effect of various PCD variants on directed ciliary particle transport has yet to be extensively analyzed through quantifiable signatures.

PCD diagnosis is challenging and often relies on several complementary approaches [8]. These include genetic testing, measurements of nasal nitride oxide (nNO), transmission electron microscopy (TEM) of ciliary ultrastructure, and high-speed video microscopy (HSVM) analysis for ciliary waveform and beat frequency (CBF) assessment. However, these approaches have inherent limitations and are technically demanding. For instance, nNO levels can be within the normal range in some individuals with PCD [9], while TEM analysis of ciliary ultrastructure can miss the diagnosis in 30% of genetically confirmed cases [10, 11]. Manual assessment of ciliary beat patterns from HSVM analysis requires significant expertise, specialized equipment and are often only performed at research centers [12, 13]. These collective diagnostic complexities and challenges highlight the need for more accessible, objective and standardized methods for accurately assessing ciliary function in PCD.

The systematization of HSVM has promoted distinct approaches for investigating cilia motility, largely divided between characterizing individual ciliary beat patterns on immersed cells and quantifying collective ciliary transport in air-liquid interface (ALI) cultures. Analyzing the beat patterns of individual, immersed fresh cilia provides data on their fundamental mechanical properties, however, the latter approach yields a more physiologically relevant assessment by capturing the emergent effects of the complete mucociliary system [14, 15]. Investigation of *in vitro* mucociliary transport is facilitated through cilia-driven particle tracking in ALI cultures. While notable prior studies have leveraged HSVM and particle tracking to compare mucociliary transport in healthy and PCD ALI cultures, these analyses have typically been restricted to few kinematic variables or limited sample sizes [15–18].

Recent efforts to standardize and automate PCD diagnostics employed machine learning approaches across various data modalities [14, 15, 19–21]. Specifically, published work on the automated classification of ciliary motion from HSVM data has relied on using a classical machine learning pipeline using Support Vector Machine that classifies pre-defined features [14], or an ensuing exploratory study applying deep learning which used a 3D Convolutional Neural Network (CNN) that directly learns visual patterns [20]. However, these analyses were limited to classifying the beat patterns of the cilia themselves, rather than directed particle transport, the primary physiological function of motile cilia. Furthermore, publicly available, open-source implementations of the predictive frameworks are absent, which prevent independent validation, widespread clinical adoption and downstream research tasks.

With the aim of addressing some of the current limitations in PCD diagnostic methods and gaining further insight into motile cilia dysfunction and impaired ciliary clearance, we created a set of simplified computational tools that assess and characterize cilia motility in cultured airway cells. These tools effectively track, analyze and classify the movement of fluorescent microsphere particles propelled by cilia in ALI epithelial cell cultures. From a cohort of 602 videos across 24 donors, we characterized the distinct kinematic profiles and trajectories of healthy versus PCD-driven particle transport. We then trained and validated classical machine learning models and a CNN to accurately predict disease status. All the analysis tools are encapsulated into CiliaTracks, a packaged pipeline integrating a comprehensive, open-source computational framework designed to provide an accessible tool for the objective analysis and diagnostic classification of ciliary motility.

## Results

### Creating a computational framework to assess and characterize cilia directed transport

To identify measurable parameters that can be used to differentiate a mucociliary transport defect related to PCD from normal cells, we developed a computational framework to track, analyze, and classify cilia-driven particle motility extracted from high-speed microscopy videos. Fluorescent microspheres were added to primary culture airway cells, and their movement was recorded (**Figure 1A**). A subsequent multi-step computational pipeline for the comparative analysis and classification of PCD and normal samples was then developed (**Figure 1B**). To increase the robustness of this analysis, PCD patients with different variants were recruited (**Table 1 and Supplemental Table S1**).

**Figure 1.**
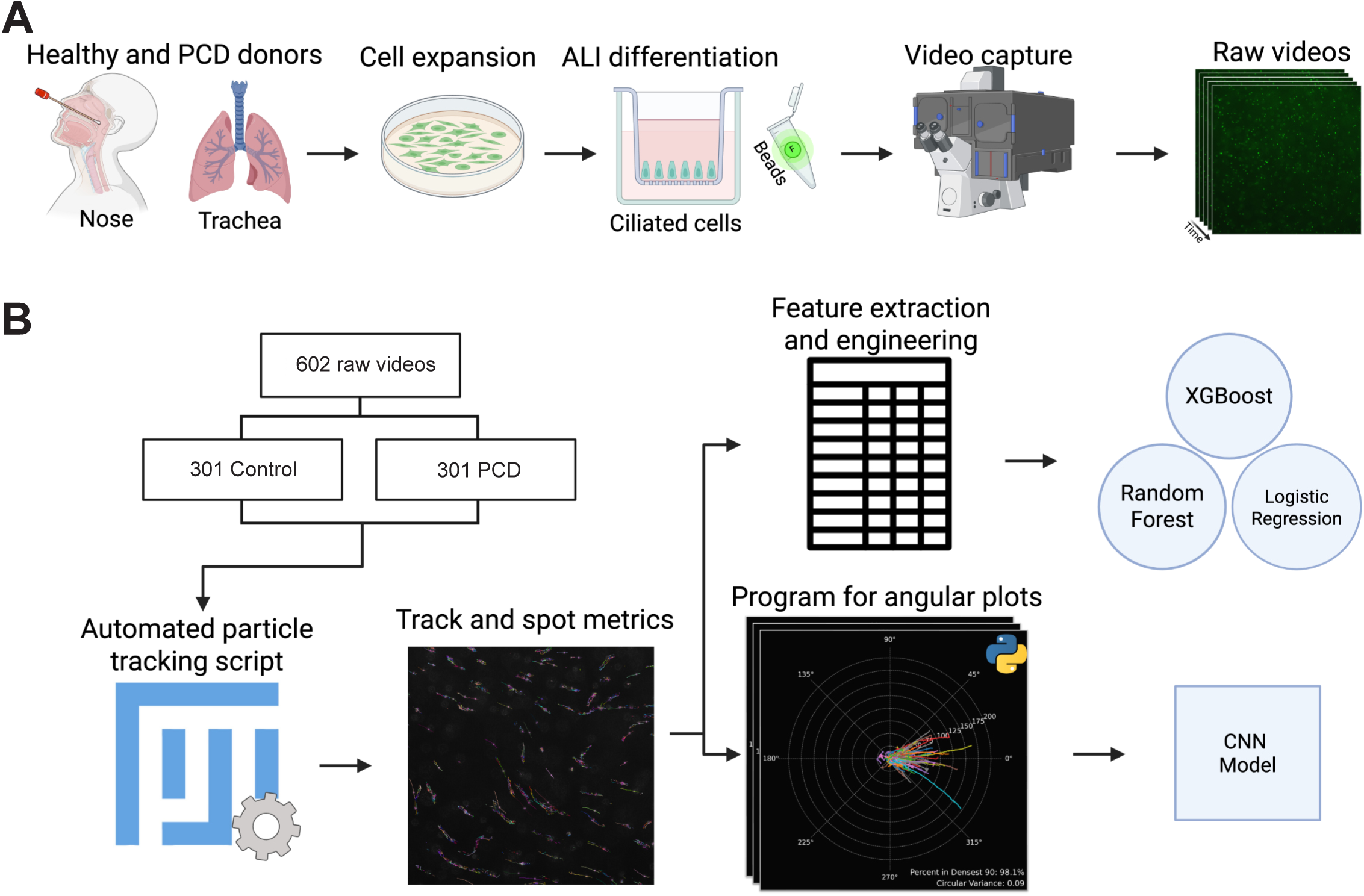
Biological and computational pipeline. **(A)** Biological workflow to acquire raw videos from cultured ciliated cells. **(B)** Schematic overview of computational approach to analyze and classify bead trajectories.

**Table 1.** Genotypes of recruited individuals.

| **Donor group** | **Gene** | **Number of donors** | **Captured videos** |
| --- | --- | --- | --- |
| Unaffected control |  | 10 | 301 |
| Primary ciliary dyskinesia | *DNAAF5* | 5 | 114 |
|  | *DNAH11* | 4 | 85 |
|  | *RSPH1* | 1 | 44 |
|  | *DNAAF1* | 1 | 21 |
|  | *DNAH5* | 1 | 13 |
|  | *HYDIN* | 1 | 12 |
|  | *RSPH4A* | 1 | 12 |

Table shows the number of donors involved in sample acquisition across PCD gene-variant and normal samples. A total of 602 videos were collected and used in the CiliaTracks analysis pipeline.

Following video acquisition, raw track metrics were extracted from recordings using an automated script developed in Fiji ImageJ. To extensively analyze these data in two parallel analytical streams, we created CiliaTracks, a custom package developed in Python. For image-based classification and visual trajectory analysis, the CiliaTracks program generated standardized angular plots. This yielded trajectory, displacement and speed polar diagrams of the cilia-propelled particles for each video. For feature-based classification and comparison, a set of 12 quantitative features that describe particle motility were extracted and quantified from the raw track data (**Table 2**). The resulting plots and feature-set data were then used to develop and validate classical machine learning models, and a Convolutional Neural Network (CNN) based model, to distinguish between healthy and PCD conditions.

**Table 2.** Track features.

| **Feature** | **Description** | **Units** |
| --- | --- | --- |
| Circular Variance | Circular spread of tracks. Ranges from 0 to 1, where 0 indicates all data points are concentrated at the mean direction, and 1 indicates a uniform distribution | Unitless |
| Confinement Ratio | Net track displacement divided by total distance | Unitless |
| Linearity of Forward Progression | Ratio between mean straight-line speed and mean speed | Unitless |
| Max Distance Travelled | Distance from origin to furthest point in the track | $\mu\text{m}$ |
| Mean Directional Change Rate | The angle between two succeeding frames, averaged over all the frames | radians/second |
| Mean Straight Line Speed | Net track displacement divided by total track time | $\mu\text{m}/\text{second}$ |
| Percent in Densest 90 | Percent of tracks in the densest 90° window | % |
| Total Distance Traveled | The full distance the particle traveled throughout a track | $\mu\text{m}$ |
| Track Displacement | Distance between first and last spot in the track | $\mu\text{m}$ |
| Track Max Speed | The maximum speed between two links in a track | $\mu\text{m}/\text{second}$ |
| Track Mean Speed | The mean speed between each succeeding link of the track | μm/second |
| Track Min Speed | The minimum speed between two links in a track | μm/second |

Track features extracted and generated from the raw output of the automated Fiji script used to process sample videos. 2 out of the 12 features were engineered to characterize global angular variance: circular variance and percent in densest 90. The rest were extracted from TrackMate's provided raw metrics.

### Trajectory plots reveal distinct PCD cilia-driven particle movement patterns

To visually compare the cilia driven particle movement patterns between healthy and PCD gene-variants, track trajectory and displacement plots were generated across all samples. Three replicates are presented for each sample group (**Figure 2 and Supplemental Figure S1**). To increase the robustness of the analysis, only the top 150 tracks sorted by mean track quality were processed. This number was selected empirically to ensure visual interpretability of the trajectory plots while minimizing noise from incomplete tracks.

**Figure 2.**
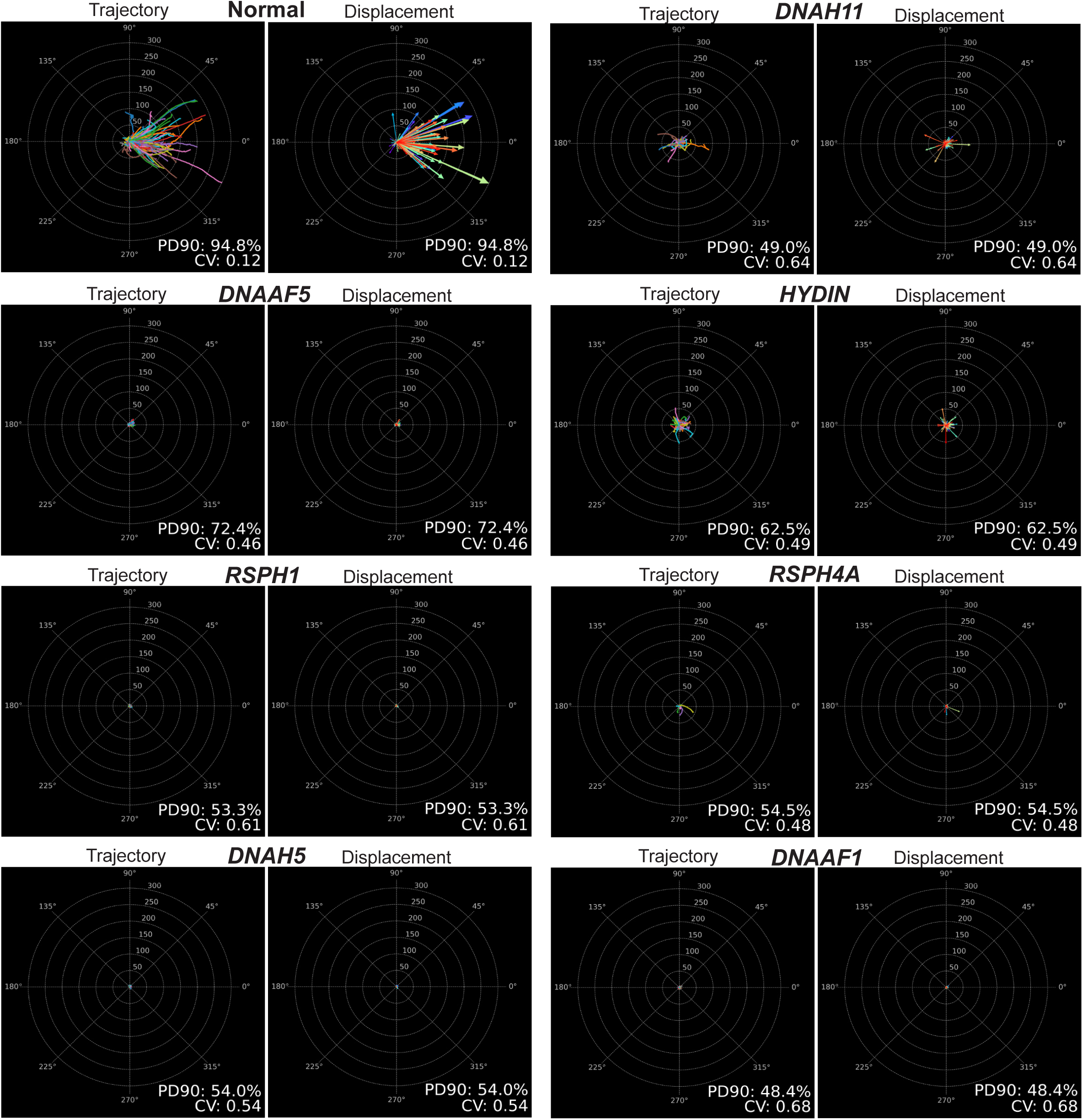
Trajectory plots showing differences in cilia-driven propulsion of microspheres between PCD and normal cells. Angular plots displaying overall particle trajectory and displacement of the top 150 tracks, normalized at an origin with standardized orientation. Select samples from the normal cohort and each genetic variant were visualized. The corresponding percent in densest 90 (represented as PD90) and circular variance (represented as CV) metrics for each sample are annotated to characterize the global angular variance of tracks.

Distinct particle movement patterns were observed among healthy and PCD cultures, coinciding with corresponding ciliary dysfunction phenotypes within the PCD samples. Normal samples typically exhibited large track displacements and aligned trajectories, with low overall angular variance. The latter was represented quantitatively by a small circular variance (CV), which is a measure of angular dispersion, with values ranging from 0 (perfect alignment) to 1 (maximum dispersion), and a high PD90, which represents the percent of tracks present in the densest 90-degree window. In contrast, PCD samples displayed reduced track displacements and disorganized trajectories with high CV and low PD90 values, often aligning with variant-specific motility defects (**Figure 2 and Supplemental Figure S1)**. Variants associated with ciliary immotility or dyskinetic motility, *DNAH5, DNAAF5*, *DNAAF1, RSPH1* and *RSPH4A* yielded plots with minimal or no discernable tracks. Conversely, variants linked to hyperkinetic motility (*DNAH11* and *HYDIN*), while displaying greater particle displacements than immotile mutants, demonstrated highly disorganized and randomly oriented trajectories, quantified by their elevated circular variance. To supplement the trajectory analysis, cilia beat frequency (CBF) in Hertz and cilia active area (% of cilia actively beating) were measured for the corresponding samples (**Supplemental Figure S2A and Supplemental Figure S2B**), using an automated software (SAVA, Ammons Engineering, MI). Genetic variants associated with hyperkinetic and dyskinetic ciliary defects (*DNAH11*, *RSPH1* and *HYDIN*) yielded elevated or similar CBF compared to the healthy samples, while PCD variants linked with ciliary immotility (*DNAAF5*, *DNAAF5*, *RSPH4A*, *DNAH5* and *DNAAF1*) displayed lower CBF. The active area was greatest in normal samples, contrasting with the limited areas of coordinated motion observed in the PCD genetic variants.

### PCD is associated with disparate movement profiles compared to normal cells

Comparative analysis of the 12 track features extracted from captured videos (**Table 2**), identified key differences in cilia-propelled microsphere movement between normal and PCD ciliated cultures. PCD samples exhibited significantly decreased track distance, displacement and speed relative to normal cells (**Figure 3A**, **Supplemental Figure S3A, and Supplemental Table S2**). Features that measure trajectory directional organization including confinement ratio (track displacement divided by total distance) and mean directional change rate (average angle between succeeding track frames), indicated a significant increase in track non-linearity and disorganization across all PCD variants (**Figure 3A**). Within the PCD cohort, motile variants *DNAH11* and *HYDIN* demonstrated the greatest speed and displacement, yet this movement was coupled with persistently elevated levels of uncoordinated trajectories. To visualize the primary axes of linear variation within the track feature dataset and assess the separability of the two cohorts, Principal component analysis (PCA) was performed, revealing a distinct separation between PCD and healthy samples along the first principal component, PC1 (**Figure 3B**). Histograms of the features further illustrated a clear distributional difference between normal and PCD samples (**Supplemental Figure S3B**). The comparative analysis indicates that PCD samples can be distinguished from normal sample videos based on a discrete set of quantifiable features.

**Figure 3.**
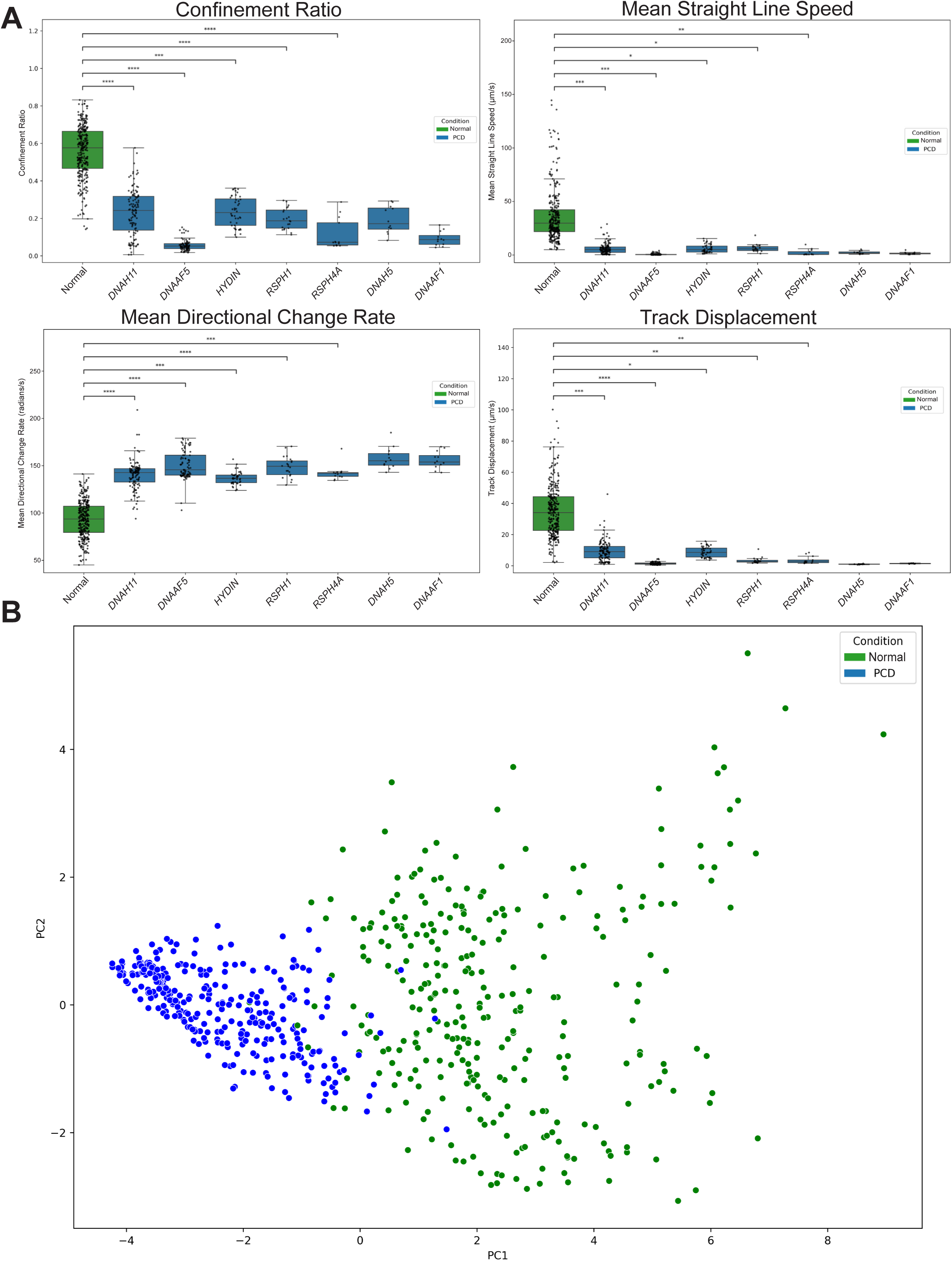
Comparison of normal and PCD track features. **(A)** Boxplots displaying select key track feature values across all individual genetic variants and normal samples (n = 32 for normal, *DNAH11* = 7, *DNAAF5* = 6, *HYDIN* = 3, *RSPH1* = 3, *RSPH4A* = 3, respectively). Groups with only 1 biological replicate (*DNAH5*, *DNAAF1*) were omitted from statistical testing. **(B)** Principal component analysis of the scaled data. The first two principal components are visualized by a scatter plot, with samples colored by condition. t-test, *p < 0.05, **p < 0.01, ***p<0.001, ****p<0.0001, ns = not significant.

### Machine learning models distinguish PCD and normal samples using track features

To objectively classify and distinguish PCD from non-PCD samples, we compared 4 machine learning models, including three classical models that were trained and tested on the 12 track features derived from the 602 captured videos, as well as a deep learning CNN model that relies on visual pattern recognition. For classical models, data was split into training (70%) and test (30%, unseen) sets, stratified by condition and PCD variant, preserving the proportional representation of each class in both sets (**Supplemental Table S3 and Figure 4A**). We employed eXtreme Gradient Boosting (XGBoost), Random Forest, and logistic regression models that were trained on features extracted from 421 samples within the training set to classify PCD and normal with a 5-fold cross validation for hyperparameter tuning (**Supplemental Table S4**). Both XGBoost and Random Forest were selected to model the complex nonlinear relationships that may exist between the features, while logistic regression was incorporated as a basic, straightforward binary classifier.

**Figure 4.**
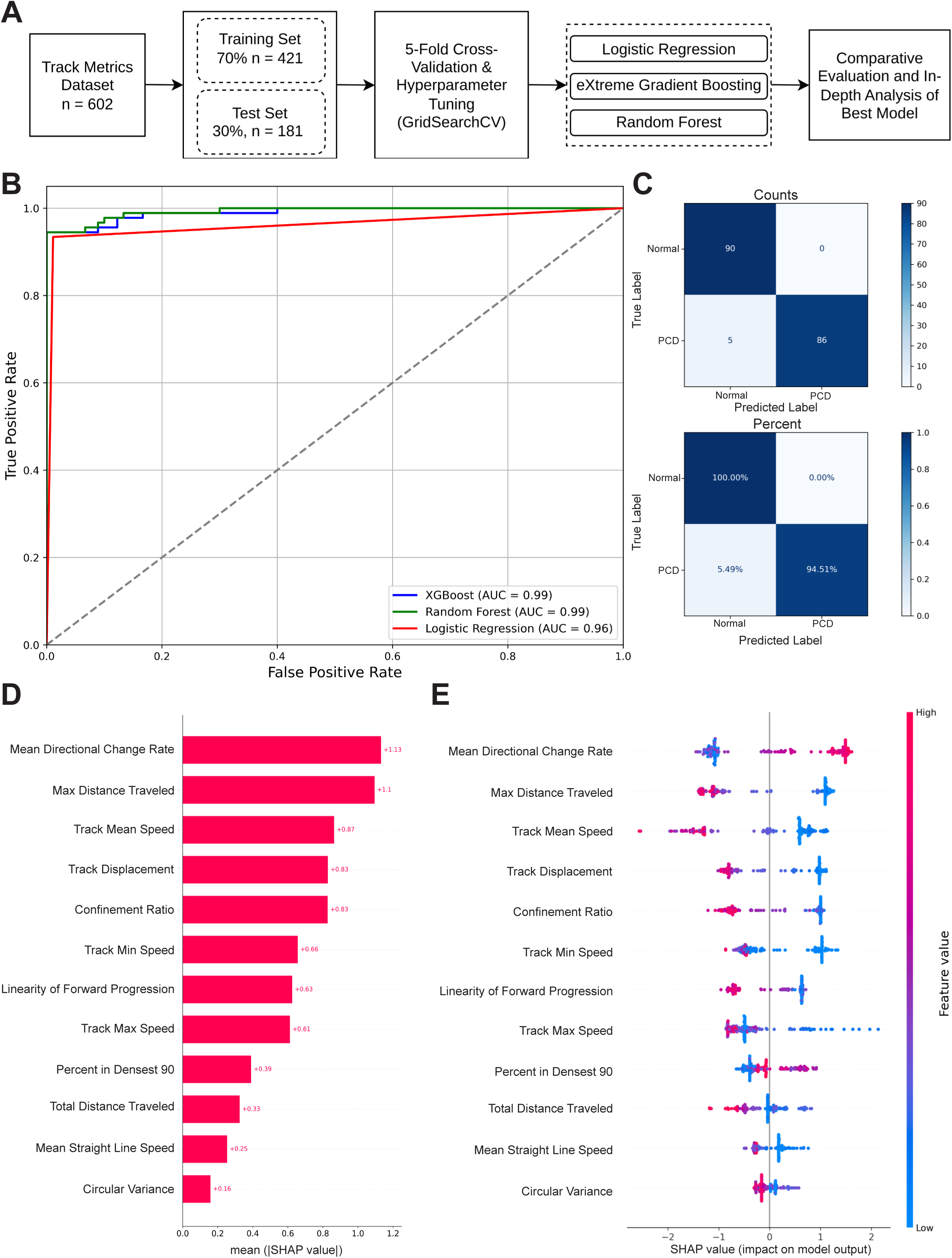
Classical machine learning model discriminates PCD from normal cells. **(A)** Flowchart showing the approach to generate and evaluate machine learning models. **(B)** Receiver operating characteristic (ROC) curves for the XGBoost, Random Forest and logistic regression classifiers on the unseen test set. The area under the curve (AUC) value for each model is displayed as a single score summarizing each model’s performance. An AUC score of 0.5 indicates null discrimination while 1.0 indicates a perfect model. **(C)** Confusion matrices for the joint-top performing classifier, XGBoost. The top panel displays the raw counts for normal and PCD sample predictions. The bottom panel shows the same result normalized by true condition. **(D)** Mean SHAP values of all 12 predictive features, illustrating the mean absolute impact of each feature on the model’s prediction. **(E)** SHAP Beeswarm summary plot for the XGBoost classifier on the test set. Each point on the plot corresponds to an individual sample. The feature’s impact on the model output for that sample is shown by its horizontal position (SHAP value). Features are ranked vertically by overall importance. The color indicates the value of the feature from low (blue) to high (red).

The ability of the models to discriminate between PCD and normal samples was evaluated on the single unseen test set (n = 181). All three classifiers demonstrated strong performance, with XGBoost and Random Forest showing excellent discriminative ability (area under the curve, AUC = 0.99 for both) (**Figure 4B**). The XGBoost and Random Forest classifiers yielded the highest accuracy and F1-scores (harmonic mean of precision and recall) on the unseen test set (**Table 3**). A confusion matrix, which tabulates the per-class counts of correct and incorrect predictions, was subsequently generated for all models (**Figure 4C and Supplemental Figure S4**), visualizing their classification accuracy. The XGBoost and Random Forest models demonstrated low misclassification rates, with only 5.5% of PCD videos incorrectly misclassified as as normal and none of normal sample videos misidentified as PCD. An overall accuracy of 0.97 was achieved for both.

**Table 3.** Prediction performance scores of classical machine learning and convolutional neural network models.

| **Model** | **Class** | **Precision** | **Recall** | **F1-Score** | **Accuracy** |
| --- | --- | --- | --- | --- | --- |
| XGBoost | Normal | 0.95 | 1.00 | 0.97 | 0.97 |
|  | PCD | 1.00 | 0.95 | 0.97 |  |
| Random Forest | Normal | 0.95 | 1.00 | 0.97 | 0.97 |
|  | PCD | 1.00 | 0.95 | 0.97 |  |
| Logistic regression | Normal | 0.94 | 0.99 | 0.96 | 0.96 |
|  | PCD | 0.99 | 0.93 | 0.96 |  |
| CNN | Normal | 0.95 | 0.96 | 0.95 | 0.95 |
|  | PCD | 0.95 | 0.95 | 0.95 |  |

Precision (proportion of items predicted as positive that are true positives), recall (ratio of true positives to the total number of actual positives), F1-score (harmonic mean of precision and recall) and accuracy were calculated for each model on the unseen test set, separated by class.

To interpret the XGBoost model’s predictions, SHapley Additive exPlanations (SHAP) [22] analysis was conducted to quantify the global and local impact of each track feature on classifications (**Figure 4D** and **4E**). The analysis identified the mean directional change rate as the most influential feature based on mean absolute SHAP value, where higher values (non-linear trajectories), strongly pushed the model’s prediction towards a PCD classification. Other features with significant predictive importance included max distance traveled, track mean speed, track displacement, and confinement ratio. For these features, the SHAP values showed a clear separation between feature magnitude and contribution (**Figure 4E**).

### Image-based classification can distinguish PCD from non-PCD samples

Since feature-based analysis may be biased by the chosen pre-conceived features, we also constructed an image-based deep learning framework using a CNN. The CNN model was directly trained and tested on the trajectory plots, to differentiate between PCD and normal samples by learning discriminative patterns directly from the images, thereby bypassing the need for manual feature extraction and selection (**Figure 5A** and **B, and Supplemental Table S5**). The CNN based model was trained on the same training set (n = 421) and evaluated on the same unseen test set (n = 181) as the classical machine learning models.

**Figure 5.**
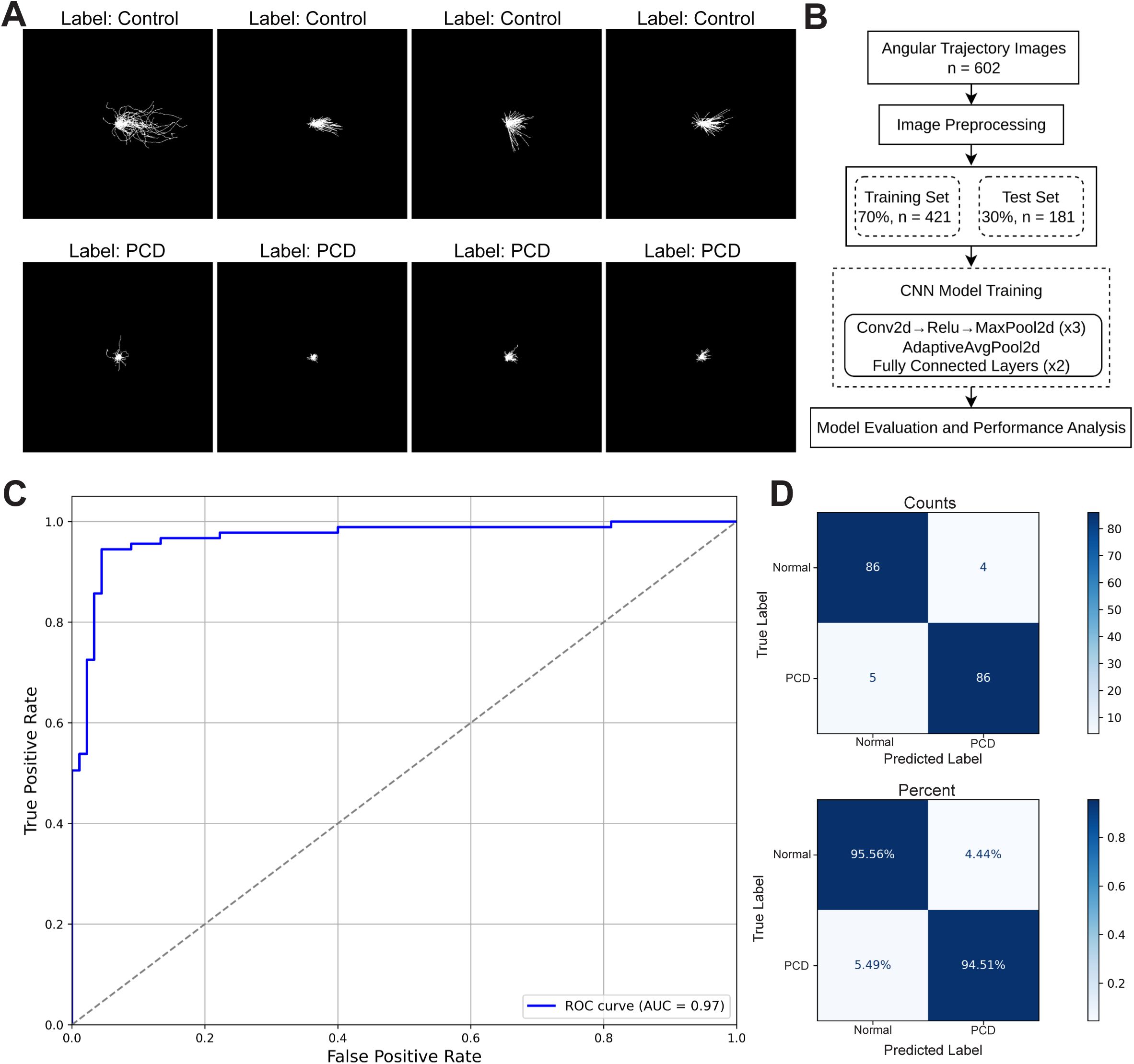
Image-based classification using convolutional neural network discriminates PCD from normal samples. **(A)** Select processed normal and PCD trajectory plots to represent example inputs for the CNN model. **(B)** CNN workflow overviewing model training and construction. (**C)** ROC curve for the CNN model predictions on the unseen test set. The AUC is annotated as a single score, where a score of 0.5 indicates null discrimination while 1.0 signifies perfect discrimination. **(D)** Confusion matrices for the CNN model. Both raw counts (top panel) and a normalized percent matrix (bottom panel) are displayed.

Evaluation of the model on the unseen test set revealed an excellent ability to discriminate between classes across varying thresholds, yielding an AUC score of 0.97 (**Figure 5C** and **5D**). The CNN achieved a strong overall accuracy of 0.95 and was effective in identifying the PCD video samples (**Table 3**). Its recall (sensitivity) of 0.95 was among the highest of the tested models, though the feature-based XGBoost and Random Forest classifiers yielded a greater overall accuracy (0.97) and superior precision (0.99) (**Table 3**). The high sensitivities are an indicator of the models’ diagnostic potential, as they correspond to a low false negative rate, with only 5.5% of true PCD samples being misclassified as normal (**Figure 5D**).

Furthermore, to validate the ability of the machine learning workflow to make predictions on new donors, the XGBoost and CNN models were retrained and tested on data that was grouped by donors (**Supplemental Table S6**). This ensured that no single donor’s data appears in both training and test set. The models achieved comparable excellent performance (**Supplemental Table S7**), demonstrating their capacity to make accurate classifications on data from unseen donors/patients.

### Using cilia beat frequency as a sole input in machine learning is less accurate that functional cilia models

Cilia beat frequency assessment has traditionally been used as a tool for distinguishing PCD from normal samples by evaluating if beat frequencies are beyond the expected normal range established by a testing research lab [23–25]. However, CBF can be within normal range in several genetic variants causative of PCD [25]. To contextualize the performance of our CiliaTracks models, we established a benchmark by training a logistic regression classifier using cilia beat frequency (CBF) and Active Area, acquired using the Sisson-Ammons Video Analysis (SAVA) system on an overlapping donor/genotype cohort (**Supplemental Table S8, Supplemental Table S9 and Figure 6A**). When evaluated on the unseen test set, the CBF-based logistic regression model demonstrated good discriminative ability yielding an AUC of 0.91 (**Figure 6B**). The model had an overall accuracy of 0.84, with balanced precision, recall and F1-scores of 0.84 for both PCD and normal classes (**Table 4 and Figure 6C**). A decision boundary plot illustrates how the model separated the two conditions based on the standardized CBF and active area values (**Figure 6D**). While the CBF-based model performed well, its predictive accuracy was lower than that of the CiliaTracks models based on functional cilia measurements. Both the feature-based XGBoost classifier (0.97 accuracy, 0.99 AUC) and the image-based CNN (0.95 accuracy, 0.97 AUC) showed superior performance. Samples with a motile cilia phenotype, especially *DNAH11*, were more likely to be misclassified in all models – though more so when using CBF benchmarking (**Supplemental Table S10 and Supplemental Figure S5**).

**Figure 6.**
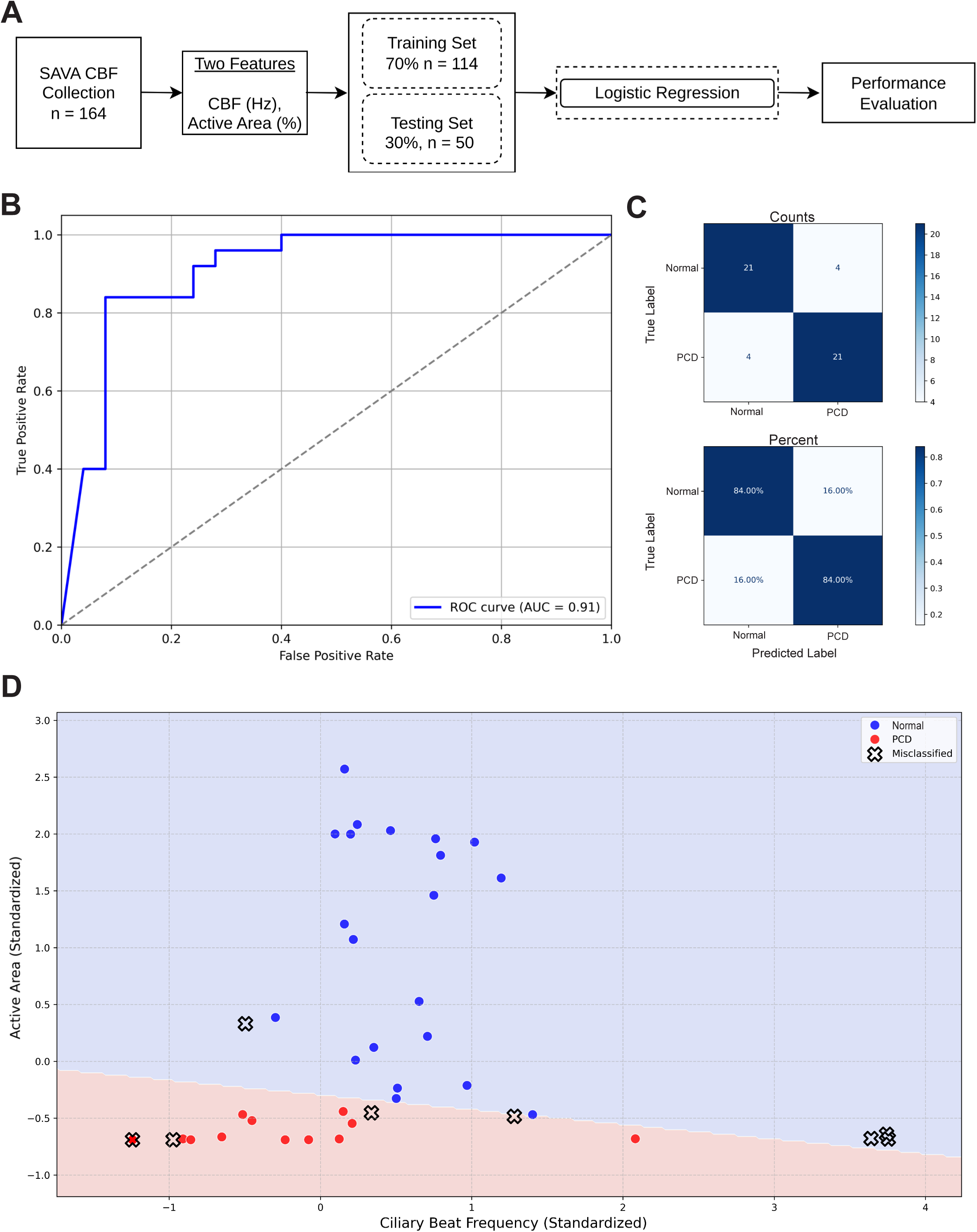
CiliaTracks shows superior performance in identifying PCD samples compared to a logistic regression model of CBF. **(A)** Workflow diagram of logistic regression model design. **(B)** ROC curve for the logistic regression model evaluated on the unseen test set. AUC score is displayed. A higher AUC score indicates a better discrimination ability. **(C)** Confusion matrices for both raw counts (top panel) and normalized percent counts (bottom panel) are shown for the model’s predictions on the test set. **(D)** Decision boundary plot of the logistic regression model on the unseen test set. Normal, PCD and misclassified data points are labelled to depict the model’s prediction accuracy. Shaded blue portion of the plot indicates standardized CBF and active area values resulting in a normal classification while shaded red portion yields a PCD classification.

**Table 4.** Cilia beat frequency logistic regression performance scores.

| **Class** | **Precision** | **Recall** | **F1-Score** | **Accuracy** |
| --- | --- | --- | --- | --- |
| Normal | 0.84 | 0.84 | 0.84 | 0.84 |
| PCD | 0.84 | 0.84 | 0.84 |  |

Table summarizes the precision, recall, F1-score and accuracy of cilia beat frequency (CBF) logistic regression model on the unseen test set for each class.

### CiliaTracks can differentiate subtypes of mucociliary defects

To evaluate whether machine learning can differentiate between different diseases affecting mucociliary function, we captured videos and analyzed samples obtained from individuals with cystic fibrosis using identical growth conditions and workflow. Cystic fibrosis is a genetic condition that disrupts ciliary clearance due to an abnormal mucus layer caused by variants in the *CFTR* gene [26]. Despite defects in clearance, angular plots of the cilia-propelled particles exhibited aligned trajectories with substantial displacements (**Figure 7A and Supplemental Figure S6**). Furthermore, comparative analysis of key kinematic track features (confinement ratio, mean straight line speed, mean directional change rate and track displacement) indicates that the cystic fibrosis tracks were characterized by elevated speed, displacement and organized trajectories compared to PCD (**Figure 7B**). Both the XGBoost and CNN based machine learning models were able to classify the cystic fibrosis samples and differentiated them from PCD samples with high accuracy (**Table 5**).

**Figure 7.**
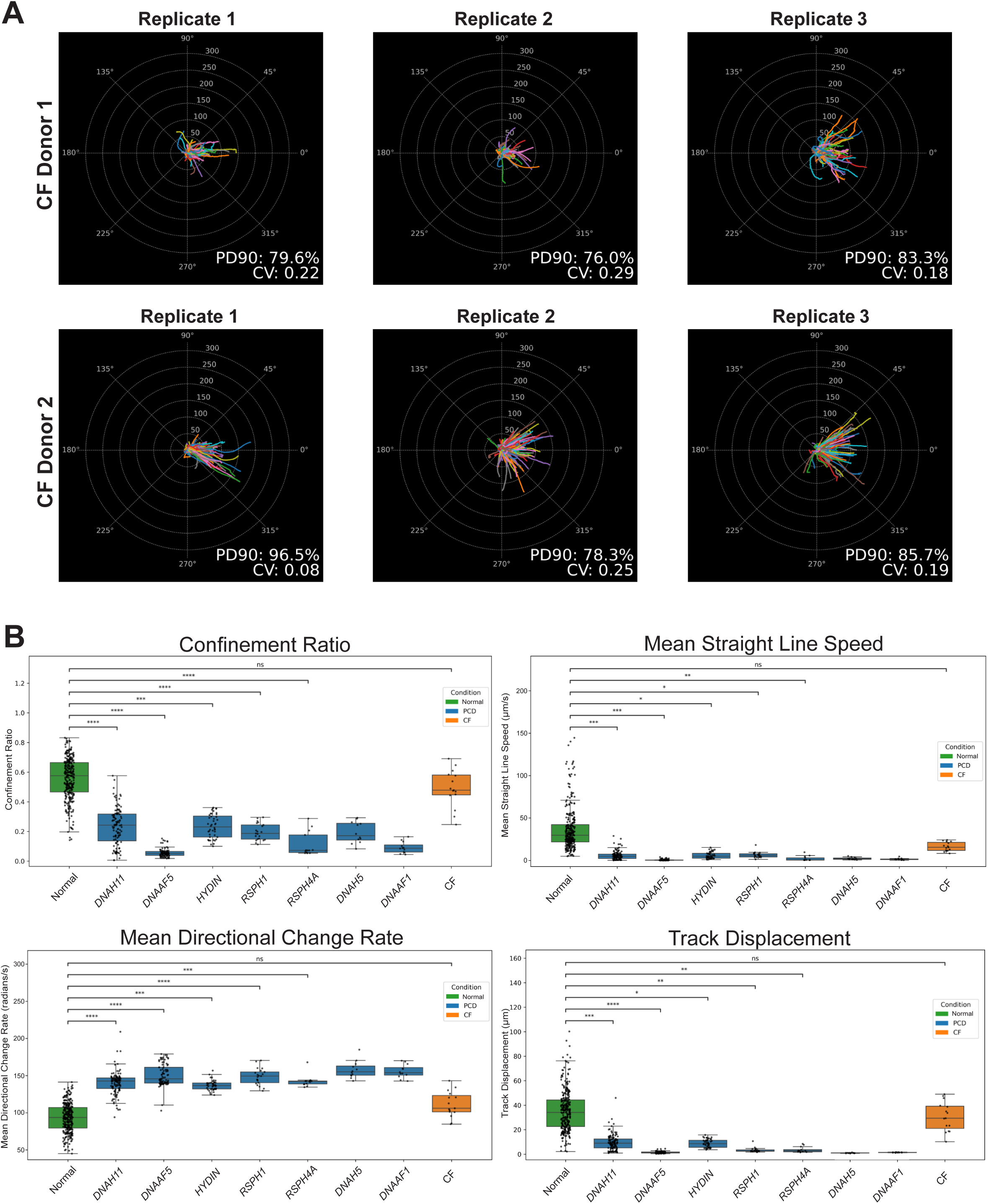
Machine learning can differentiate PCD from cystic fibrosis samples. **(A)** Trajectory and displacement angular plots of the top 150 tracks for selected cystic fibrosis videos from both donors. Tracks are normalized at an origin and standardized for direction. Circular variance (CV) and percent in densest 90 (PD90) values are annotated on the plots to characterize the global angular variance of tracks. **(B)** Box plots depicting the key track metric values for all normal, PCD and cystic fibrosis videos. PCD samples are split by genetic variant. (n= 32 for normal cells, *DNAH11* = 7, *DNAAF5* = 6, *HYDIN* = 3, *RSPH1* = 3, *RSPH4A* = 3, Cystic Fibrosis = 5, respectively; t test: *p < 0.05, **p < 0.01, ***p<0.001, ****p<0.0001, ns = not significant). Groups with only 1 biological replicate (*DNAH5*, *DNAAF1*) were omitted from statistical testing.

**Table 5.**
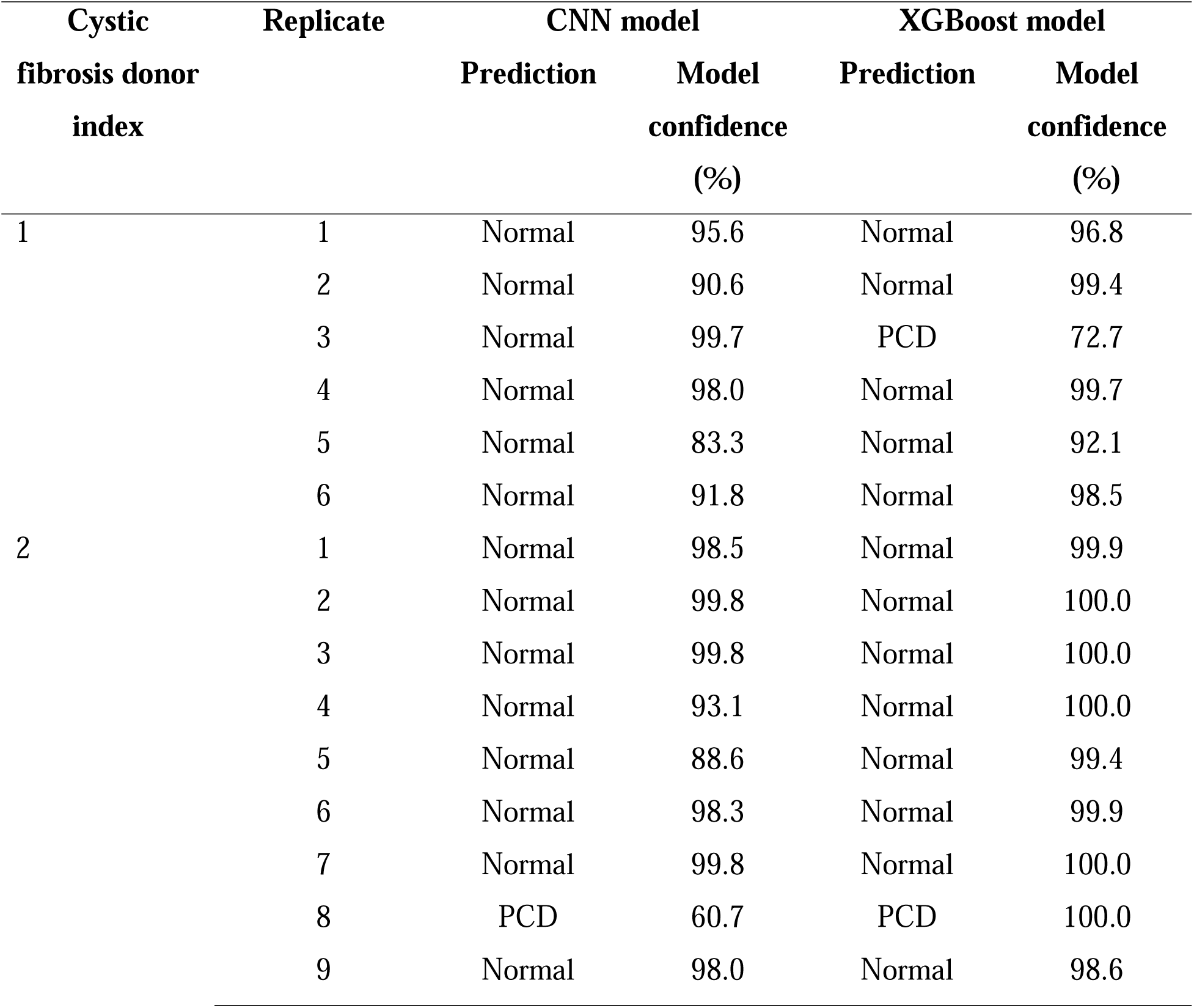
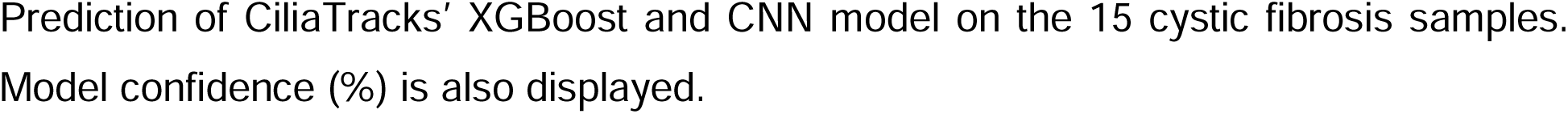
XGBoost and CNN model predictions for cystic fibrosis samples.

### Access to advanced diagnostic tools

To facilitate access to advanced diagnostic framework, we created a desktop application and Python package to ensure the accessibility of the presented tools. The application and package host all the CiliaTracks functions and enables users to seamlessly upload raw TrackMate outputs CSVs to obtain a prediction from both XGBoost and CNN models. The application works across Linux, Windows and MacOS. While the Python package can be installed from the Python Package Index (PyPI), under CiliaTracks. The complete source code, documentation, and the automated Jython TrackMate script will be made publicly available. To promote greater accuracy and generalizability, the publicly deployed XGBoost and CNN models were trained on all the available data. These models will be periodically retrained and updated as new data becomes available to enhance their performance over time.

## Discussion

The analysis of cilia-propelled particle movement produces a proxy and physiologically relevant measure of ciliary clearance, the primary function of motile cilia in human airways. Here we attempted to provide a large and comprehensive quantitative analysis of cilia-driven transport, revealing distinct particle movement profiles between healthy and PCD samples. Here we developed a comprehensive, open-source computational framework designed to objectively analyze and classify cilia-driven particle motility from culture multiciliated cells. This quantitative analysis of ciliary transport reveals a consistent and measurable signature of ciliary function that effectively differentiates healthy and diseased cultures. The comparative characterization of particle transport using trajectory patterns and kinematic track features revealed a persistent separation between normal and PCD cohorts.

Normal ciliary transport is typically categorized by substantial coordination and uniformity resulting in high speed, aligned trajectories with minimal angular variation [6, 7]. Our analysis of PCD captured videos from different variants showed a clear breakdown of this organized movement. For individuals with variants leading to immotile or dyskinetic cilia (*DNAH5, DNAAF5*, *DNAAF1, RSPH1* and *RSPH4A*), trajectory plots displayed minimal or no discernible displacement, reflecting a failure of the ciliary clearance mechanism (**Figure 2 and Supplemental Figure S1**). Meanwhile, variants linked to hyperkinetic beating (*DNAH11* and *HYDIN*), exhibited disorganized and erratic trajectories, with significant angular dispersion. This erratic motion is consistent with the established understanding of how ciliary disorientation and discoordination impair the propulsive beating necessary for effective fluid transport, even when beat frequency is in the normal or hyperkinetic range [27].

The disparity of cilia movement patterns was further defined by the analysis of 12 quantitative track features, showing key kinematic differences between healthy and PCD cohorts (**Figure 3A and Supplemental Figure 3A**). Principal component analysis based on these features demonstrated a distinct linear separation between the conditions (**Figure 3B**). Features describing non-uniform trajectory motion, including confinement ratio and mean directional change rate, were significantly elevated in PCD samples, highlighting the effects of PCD variants on cilia function beyond its frequency and speed (**Figure 3A**). This multi-faceted comparative analysis of cilia-driven particle transport provides an objective signature of cilia function and ciliary clearance, distinguishing healthy and diseased states, and could be used to overcome the subjectivity and technical limitations of existing PCD diagnostic methods.

The feature and image-based machine learning classification pipelines achieved excellent performance in differentiating disease states, showcasing that measured cilia-propelled particle movement is a reliable indicator of disease (**Table 3**). These findings are consistent with a growing body of research that shows the utility of deep learning methods as potential diagnostic modalities for PCD [19, 21].

Although machine learning based on functional assessment of beads transport was able to separate between PCD and normal conditions, there were 5 misclassified false negative video samples from the XGBoost model and 5 misclassified false negative plots from the CNN test set evaluation, all of which were due to *DNAH11* variants (**Supplemental Table S10 and Supplemental Figure S5**). This highlights the unique diagnostic challenges of this specific genotype. Variants in *DNAH11* are known to cause a subtle, hyperkinetic beating pattern with normal ciliary ultrastructure, making them difficult to diagnose by conventional methods, including HSVM (34,35). While the XGBoost and CNN model still identified 85.3% of DNAH11 samples correctly, DNAH11 cases may still require an expanded training dataset or additional diagnostic modalities to help identify subtle changes more effectively.

Since cilia beat frequency assessment is widely used as a tool to differentiate PCD from healthy samples, we constructed a CBF-based logistic regression model to benchmark CiliaTracks. The superior performance of CiliaTracks models highlights the diagnostic advantages of analyzing collective particle transport rather than traditional ciliary metrics. This was directly evidenced by the benchmark’s reduced performance on diagnostically challenging variants like *DNAH11* and *HYDIN*, where it misidentified over 50% of both variants. The disorganized, high frequency beating characteristics of these variants, which registers as ‘active’ by CBF methods, was more accurately classified by the CiliaTracks deep machine framework. The latter was able to better quantify the erratic nature of particle paths through visual trajectories and kinematic features, making it a more suitable tool for assessing ciliary function in hyperkinetic variants. While this comparison is informative, we acknowledge that it is not perfectly matched as we trained the CBF model on a smaller dataset with a different distribution of PCD variant samples. Nevertheless, the significant performance gap strongly suggests that the direct assessment of cilia-propelled particle movement is more suited for PCD diagnosis.

We also explored the framework’s evaluation of cystic fibrosis, another condition affecting mucociliary function. Although both PCD and cystic fibrosis are genetic diseases characterized by impaired mucociliary clearance (MCC), their underlying pathologies are distinct. PCD is a disorder of the intrinsic ciliary machinery, while cystic fibrosis impairs clearance due to an abnormally thick mucus layer, with the cilia remaining functionally intact [28]. In our *in vitro* system, the intrinsically normal cystic fibrosis cilia generated transport patterns that largely resembled those of healthy controls, evidenced by our model classifications (**Table 5**). This ability to differentiate between distinct mechanisms of mucociliary failure showcases the transferrable potential of CiliaTracks to investigate a variety of respiratory conditions.

The overall dataset was equally partitioned between healthy and PCD samples, however, it is worth noting that the PCD cohort itself is imbalanced across genotypes, evidenced by a differing number of samples and donors. When variants are grouped by their motility phenotype (immotile vs hyperkinetic/dyskinetic), the dataset becomes reasonably balanced, potentially mitigating model learning bias. For those reasons, although our models are robust for classifying disease and healthy samples, they have not been validated to predict individual pathogenic variants. Nonetheless, we intend to explore CiliaTracks’ predictive performance on individual variants after further diversifying the PCD genotype and expanding the data in which the models are trained on.

Air-liquid interface cultures remain the nearest approximation to the complex physiological *in vivo* environment of human airways [29]. Nonetheless, they still have inherent limitations. They do not replicate mucus rheology, immune cell presence and the mechanical forces of breathing *in vivo* [6]. This acknowledges that while an *in vitro* methodology of transport is highly relevant, the measured particle transport serves as an incomplete model for the more complex process of mucociliary clearance *in vivo*. Furthermore, it is important to consider that based on established physical principles, the residual movement of microspheres in samples with immotile cilia (*DNAH5, DNAAF5*, *DNAAF1*) is expected to be governed by Brownian motion. Therefore, it is possible that the model’s predictions are based not only on classifying cilia function but also on distinguishing active biological transport from the consistent patterns of passive diffusion.

Both feature-based and image-based models provide unique and valuable contributions to a diagnostic workflow. The interpretable nature of the XGBoost model provides insights into the specific kinematic variables driving cilia transport dysfunction, while a convolutional neural network (CNN) model provides a general, automated solution that is suited for rapid screening. By providing both tools in a publicly accessible framework, we envision researchers and clinicians adopting the high-throughput analysis of mucociliary transport as a diagnostic tool for PCD. The tools provided by the CiliaTracks will allow others to generate input data for their own classifiers, fine-tune the CNN model and investigate the disparate complexities of ciliary transport across different PCD genotypes.

Ultimately, this work establishes the quantitative analysis of cilia-driven particle transport as a methodology for PCD classification and provides an accessible, open-source framework to implement this approach in both clinical and research settings.

## Methods

### Patients

Following a protocol authorized by the Washington University Institutional Review Board (IRB# 201705095). Participants were recruited from the PCD and Rare Airway Disease clinic at St. Louis Children’s Hospital and Washington University. These included individuals with a diagnosis of PCD confirmed by genetic testing, their healthy family members, and individuals with a confirmed diagnosis of cystic fibrosis. Nasal cells were retrieved from the inferior nasal turbinate using a cytology brush, while cells retrieved from excess tissue from lungs donated from transplantations were used as additional controls as previously described [30].

### Sample preparation

Primary airway cells were processed and grown in culture using air-liquid phase conditions (ALI) as previously described [29, 31]. Samples were collected from the 24 donors in (**Table 1**). The PCD cohort was comprised of 14 individuals with seven distinct PCD genetic variants including *DNAAF5*, *DNAH11*, *RSPH1*, *DNAAF1*, *DNAH5*, *HYDIN* and *RSPH4A*. Tracheal epithelial cells samples were obtained from 10 healthy individuals. While an additional two cystic fibrosis donors homozygous for the CFTR p.F508del variant were processed identically and used for downstream analysis.

### Imaging

Cultured cells were first washed with warmed media to remove debris and dead cells. 1.9 mm fluorescent polymer microspheres (Thermo-Fisher Scientific) were diluted 1:1000 using 2% Nu-Serum (Corning) and added to the upper chamber of the ALI cultures. The volume was adjusted to the size of the culture Transwell (Corning) transwell, with a total of 20 µL used for 6.5 mm diameter transwells and 60 μL for 10mm transwells. Cells were then incubated for 10 minutes at 37 °C before imaging. Fluorescent microspheres movement was imaged live and recorded using an inverted Nikon Ti-E microscope housed in an environmental chamber maintained at 37 °C. High resolution 4.45 second videos were captured using 10x objective lens at 100 frames per second, totaling 445 frames per file. 602 videos were captured across the unaffected controls and PCD cells (**Table 1**). A minimum of three videos were recorded from random regions on each Transwell insert.

### Automated particle tracking

A custom Jython script was developed in Fiji ImageJ software (Wayne Rasband, NIH v1.54p) using the TrackMate plugin (v17.12.0) to automate the tracking of fluorescent microspheres and the subsequent extraction of track metrics (e.g. distance and speed) [32, 33]. The code was optimized to handle the processing of multiple videos iteratively. The script initially processes videos by flipping the depth and time dimensions and converting them to grayscale from RGB. For the detection of individual particles, a Laplacian of Gausian (LoG) detector is used, set with a minimum radius of 3.5 pixels. A classical quality filter of 0 is then applied using the FeatureFilter function. Next, to track the particles, a Kalman filter-based tracker is called by the script using an initial search radius of 20 pixels and maximum frame gap of 1. The track and spot metrics are subsequently saved as CSV files. The script is available for download on our CiliaTracks GitHub page.

### Trajectory visualization

Track trajectory and displacement plots were generated for each video sample using a custom-built CiliaTracks Python (v3.11) package based off the raw track metrics produced by the automated particle tracking. CiliaTracks also enables the visualization of polar speed plots. Matplotlib (v3.8.4), pandas (v2.1.4) and numpy (v1.26.4) were used in the development of these functions. When designing the functions, only top 150 tracks of each sample were considered. These were selected by mean track quality, a metric calculated by Fiji’s TrackMate plugin which incorporates the mean, max, median and standard deviation of all spots in a track. Plotting of the top 150 tracks maintains the visual interpretability of the trajectories and minimizes the noise from incomplete tracks. To standardize the overall track orientations, all trajectories were rotated by subtracting a central orientation angle (θ_mean_). This central angle was calculated using only the subset of tracks with a net displacement greater than the sample’s mean displacement. The angle of each of these *n* tracks (θ_i_) was first determined from the change in its *x* and *y* positions (Δx,Δy), between the start and end of the track:

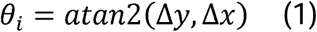

The central orientation angle (θ_mean_) was then calculated as the angle of the mean resultant vector of these tracks:

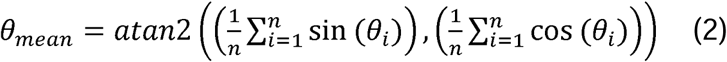

Two features were engineered to describe overall angular variance of the tracks within a sample: percent densest in 90° (PD90) and circular variance (CV). The former was found by identifying the 90° window containing the maximum percentage of track angles and calculating the overall percent of tracks within this window. Circular variance, a direct measure of angular dispersion, was calculated from the length of the mean resultant vector, *R*:

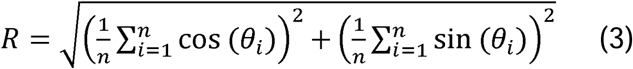

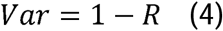

### Track feature analysis

To explore the kinematic properties of the tracks and approximate a quantifiable signature for mucociliary transport, we extracted 10 and engineered 2 (PCD90, CV) features from the automated Fiji script’s raw outputs (**Table 2**). Collectively, these 12 features measure the track speed (track max speed, track mean speed, track min speed, mean straight line speed), distance (total distance traveled, max distance traveled), displacement (track displacement), linearity (linearity of forward progression, mean directional change rate, confinement ratio) and the global angular variance (circular variance, percent in densest 90). All features derived from 602 input videos were concatenated into a single structured dataset (602 x 12) using pandas (v2.1.4) for downstream comparative analysis. To assess the variation in the dataset and explore feature contributions, a principal component analysis (PCA) was performed on a scaled version of the data using scikit-learn (v1.5.1) (https://www.jmlr.org/papers/volume12/pedregosa11a/pedregosa11a.pdf). The first two principal components were visualized in a scatter plot, with samples colored according to condition.

### Cilia beat frequency analysis

To complement the trajectory analysis of the samples, Cilia Beat Frequency (CBF) analysis was performed as previously described [34]. Images were captured using a Nikon Eclipse Ti-U inverted microscope housed in a 37°C chamber and a high-speed video imaging (CMOS camera) and Sisson-Ammons Video Analysis system processing (Ammons Engineering) [35]. CBF and Active Area (% area with active cilia beating) were analyzed in a minimum of three fields for each relevant donor (**Supplemental Figure 2).**

### Classical machine learning for track feature-based disease classification

Three different classification algorithms were developed: an XGBoost (eXtreme Gradient Boosting) classifier (v2.1.1) [36], and a Random Forest and logistic regression model from the scikit-learn library (v1.5.1). XGBoost and Random Forest are ensemble tree-based classifiers that learn complex nonlinear relationships between features through multiple decision trees, while logistic regression is a simple linear model that estimates class probabilities through a weighted combination of features (18, https://www.jmlr.org/papers/volume12/pedregosa11a/pedregosa11a.pdf). The 602 x 12 dataset was split into a training (70%, n = 421) and a test set (30%, n =181) that was left unseen for unbiased evaluation (**Supplemental Table S3**). This split was stratified by both condition (normal, PCD) and by genetic variant to promote generalizability. Hyperparameter optimization was performed on the three classifiers, which were tuned using a 5-fold cross-validated grid search implemented with the GridSearchCV function in scikit-learn only on the training data. Prior to optimization, features for the logistic regression model were standardized. The models were then instantiated with their best hyperparameters (**Supplement Table S4**), and performance was assessed on the unseen test set using typical evaluation metrics including accuracy, precision, recall and F1-score (harmonic mean of precision and recall). The models’ ability to discriminate between classes was evaluated using the Receiver Operating Characteristic (ROC) curve and the corresponding area under the curve (AUC) score. The ROC curve illustrates the discriminative ability of the binary classifiers by plotting the true positive rate against the false positive rate. The AUC score then summarizes the overall plot, where a higher value indicates better discriminative performance. A per-video evaluation framework was adopted to provide a granular and stringent assessment of performance on localized motility patterns. SHapley Additive exPlanations (SHAP) analysis (v0.46.0) was conducted on the best-performing classifier, XGBoost, to illustrate the impact and magnitude of each feature on the model’s output for the unseen test set (https://arxiv.org/abs/1705.07874). This approach enables both global interpretation of overall feature importance and local interpretation of how individual features drive specific predictions.

### Convolutional Neural Network for trajectory plot-based disease classification

Using our CiliaTracks Python package, the videos were processed into 602 grayscale trajectory plots for analysis. Each grayscale image was resized to 500×500 pixels. Image pixel values were normalized to a range of [0, 1], and the corresponding condition labels were numerically encoded. The entire dataset was then converted into PyTorch (v2.3.1) tensors. For model training and validation, the dataset was divided into training and testing sets using a stratified 70/30 split, matching the sample identities from the classical machine learning workflow. A sequential CNN was constructed in PyTorch to perform binary classification. The architecture (**Supplemental Table S5**), is composed of three convolutional blocks, followed by a classifier head. Each convolutional block contains a 2D convolution layer with a 3×3 kernel, a Rectified Linear Unit (ReLU) activation function, and a 2×2 max-pooling layer (stride 2) for spatial down sampling [37]. The blocks progressively increase feature map depth from 16 to 64.

The classifier head consists of a global adaptive pooling layer, which condenses each feature map into a single value, producing a fixed-size feature vector. This vector is then processed by two fully connected (dense) layers: a hidden layer with 128 neurons (ReLU activated) and a final output layer with 2 neurons yielding raw logits for classification.

The model was trained for 15 epochs on a system equipped with CUDA (v12.1). The Adam optimizer [38] was used for weight updates with a learning rate of 0.001 and a batch size of 4. The cross-entropy loss function was employed to handle the binary classification task. Model performance was assessed on the held-out test set. The scikit-learn library was used to generate precision, recall and F1-scores along with a confusion matrix and ROC curve for performance assessment.

### Model training and validation grouped by donor

To ensure that the performance of the models was not biased by potential donor data leakage between the training and tests sets, the models were retrained and evaluated using a group-based approach. The StratifiedGroupKFold from scikit-learn (v.1.5.1) was used to partition the data by grouping all samples from each unique donor identity and further partitioned by condition (PCD, Normal) (**Supplemental Table S6**). The XGBoost and CNN model were then retrained and evaluated, demonstrating similar strong performance (**Supplemental Table S7**).

### Benchmarking CiliaTracks models with CBF assessment

To benchmark the performance of our CiliaTracks models with an established tool for ciliary analysis, we used Sisson-Ammons Video Analysis (SAVA) system processing (Ammons engineering) CBF assessment (27). CBF analysis was carried out on samples from matching genotypes and donors used to train our CiliaTracks models, as previously described (26). The cultures were analyzed using a Nikon Eclipse Ti-U inverted microscope housed in a 37°C chamber and a high-speed imaging CMOS camera. CBF (Hertz) and Active Area (% area with active cilia beating) were analyzed in a minimum of 3 fields obtained for each relevant donor. The final dataset totaled 164 captured data points across normal and PCD cultured cells (**Supplemental Table S8**). A logistic regression classifier was then trained and evaluated on the two features, CBF and Active Area using a 70% training and 30% test set (**Supplemental Table S9**) to distinguish PCD and normal samples. The training and test set were stratified by condition and genotype, aligning with CiliaTracks models. The performance of this benchmark classifier was then assessed on the unseen test using the same evaluation metrics.

### Cystic fibrosis analysis

Samples from two donors with confirmed diagnosis of cystic fibrosis were acquired to investigate the cilia-propelled movement of a non-PCD condition that also impairs mucociliary clearance (MCC). The samples were processed using the identical biological and computational workflow as the PCD and normal cohort, to ensure methodological consistency. Using the CiliaTracks pipeline, we generated angular plots of particle movement and compared key track features with the existing dataset. Following this analysis, our XGBoost and CNN models, trained on all available, were used to classify the cystic fibrosis videos.

## Acknowledgements

The authors would like to thank Dr. Benjamin Gaston and Michael D Davis from the division of Pulmonology, Allergy, and Sleep Medicine at Riley Hospital for Children at Indiana University School of Medicine for contributing material used in this work.

