## Supplemental material for "Machine Learning Analysis of Cilia-Driven Particle Transport Distinguishes Primary Ciliary Dyskinesia Cilia from Normal Cilia"

Conflict of interest statement: All authors declare that no conflict of interest exists.

**Short title: Deep Learning of Particle Transport Identifies PCD**

**Corresponding author:**

Amjad Horani, MD

Department of Pediatrics, 660 South Euclid Avenue, Mailbox 8116. St. Louis, Missouri, 63110.

ORCID: 0000-0002-5352-1948

**Supplemental Figure**

**Figure S1. Visual trajectory analysis of selected samples in triplicate.**

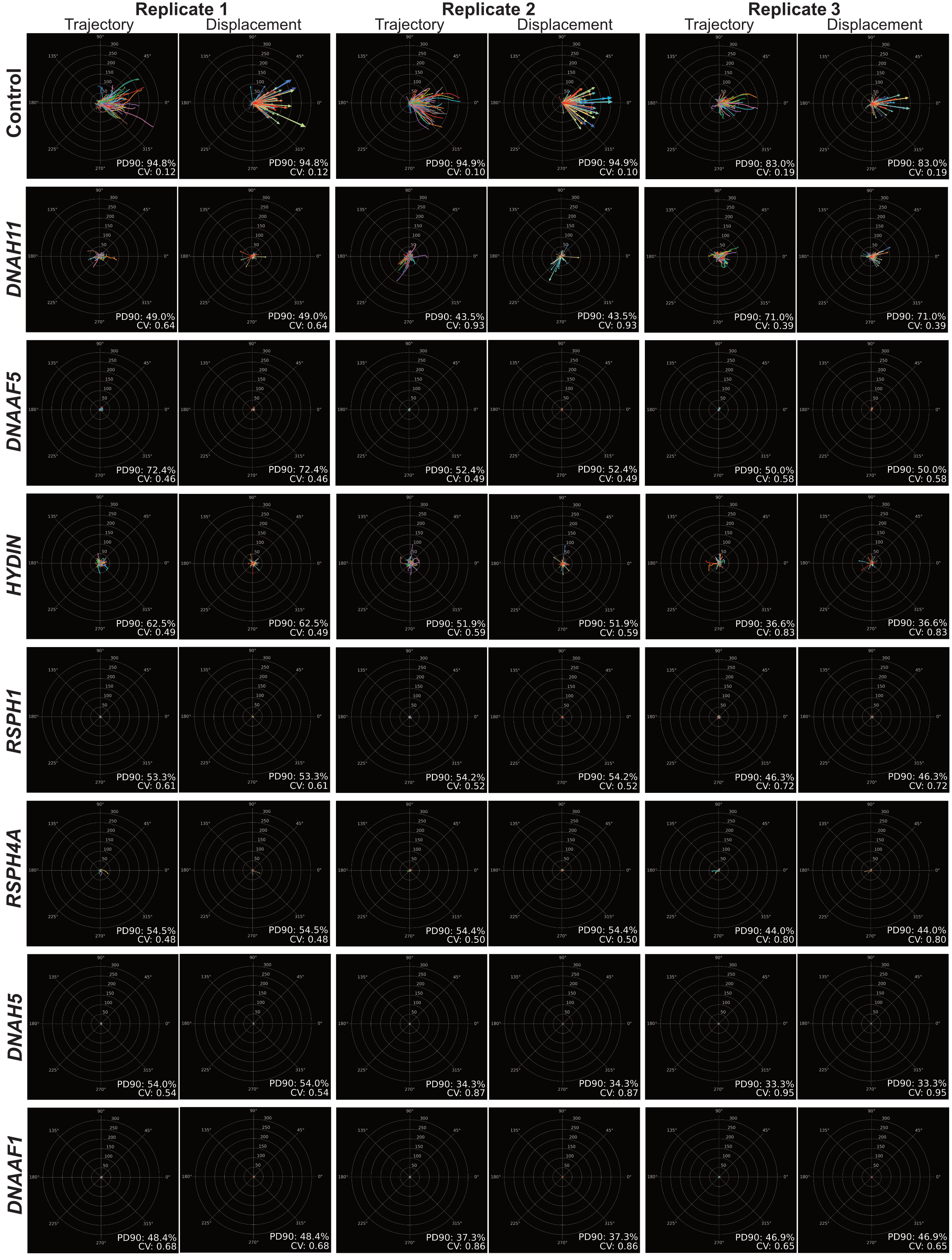

Trajectory and displacement plots of all selected video samples in triplicate from normal cells and the total PCD-variant cohort. Percent densest in 90 (PD90) and circular variance (CV) are annotated to represent the angular variance of tracks.

**Supplemental Figure S2. Cilia beat frequency analysis of PCD variants.**

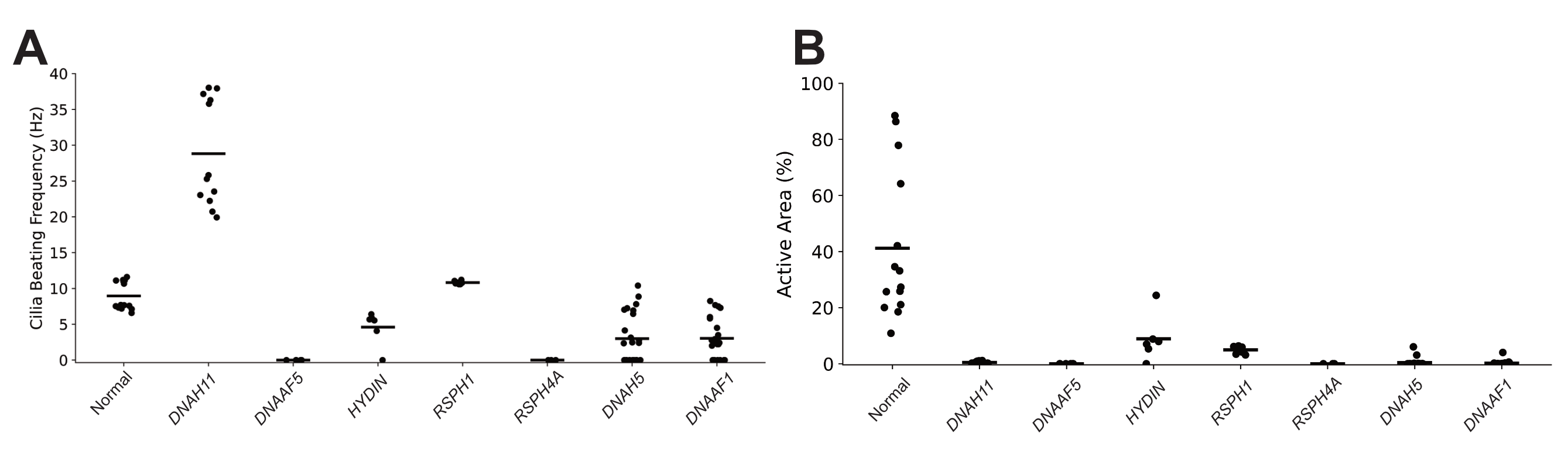

**(A)** CBF in hertz for the genetic variants and normal in shown in **Figure 2**. **(B)** The active area in % for the matching samples in **A**.

**Supplemental Figure S3. Complete statistical comparison of track feature values.**

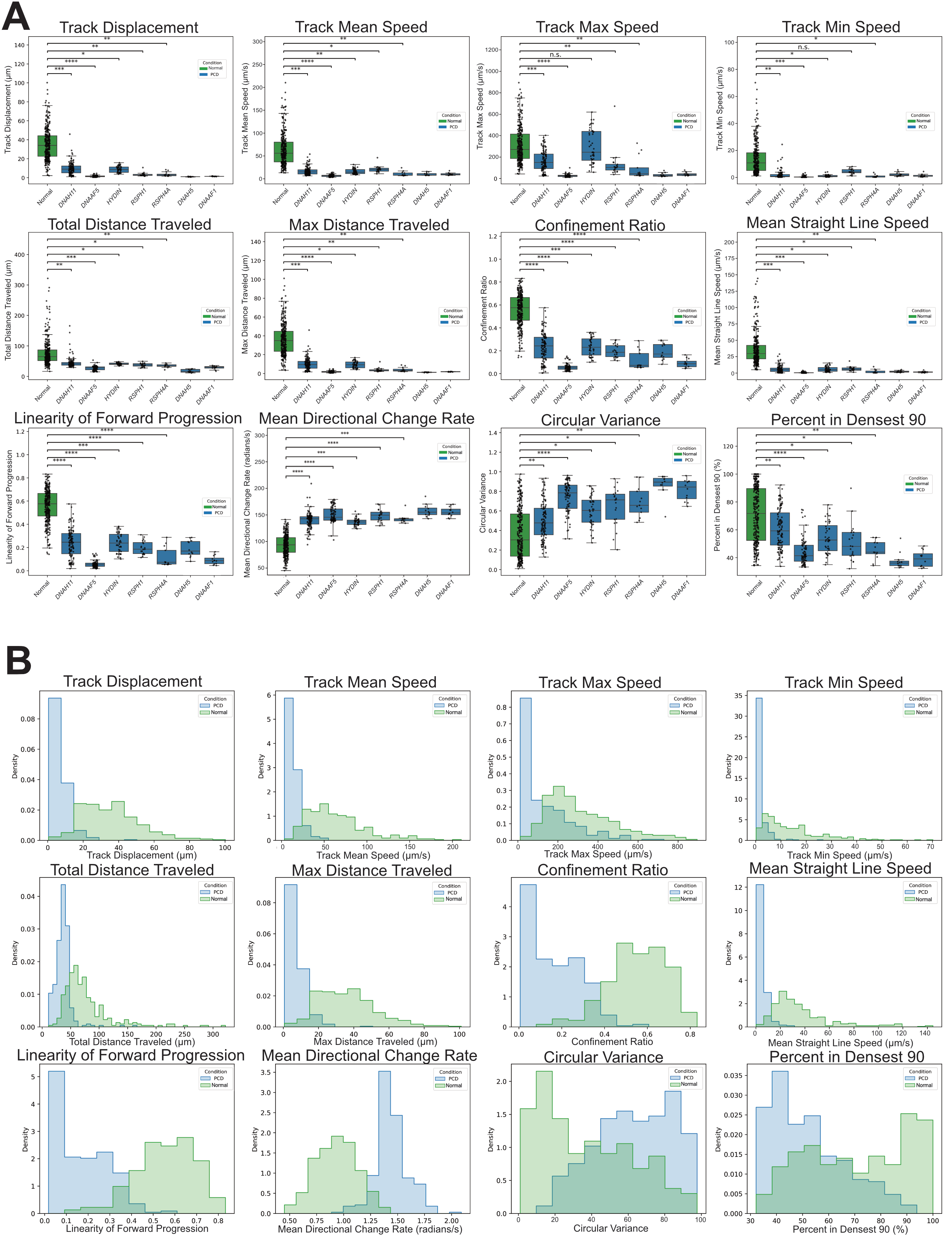

**(A)** Boxplots depicting all 12 track feature data values across individual genetic variants and normal samples. An independent samples t-test was performed to compare between normal and PCD variants, with asterisks indicating p-value significance levels. n = number of biological replicates per group (Normal: n = 32, *DNAH11*: n = 7, *DNAAF5*: n = 6, *HYDIN*: n = 3, *RSPH1*: n = 3, *RSPH4A*: n= 3), t test: *p < 0.05, **p < 0.01, ***p<0.001, ****p<0.0001, ns = not significant. Groups with only 1 biological replicate (*DNAH5*, *DNAAF1*) were omitted from statistical testing. **(B).** Density distributions of track features between conditions. Histograms depicting the density distribution of all track feature values across PCD and control samples.

**Supplemental Figure S4. Second-tier feature-based model performance.**

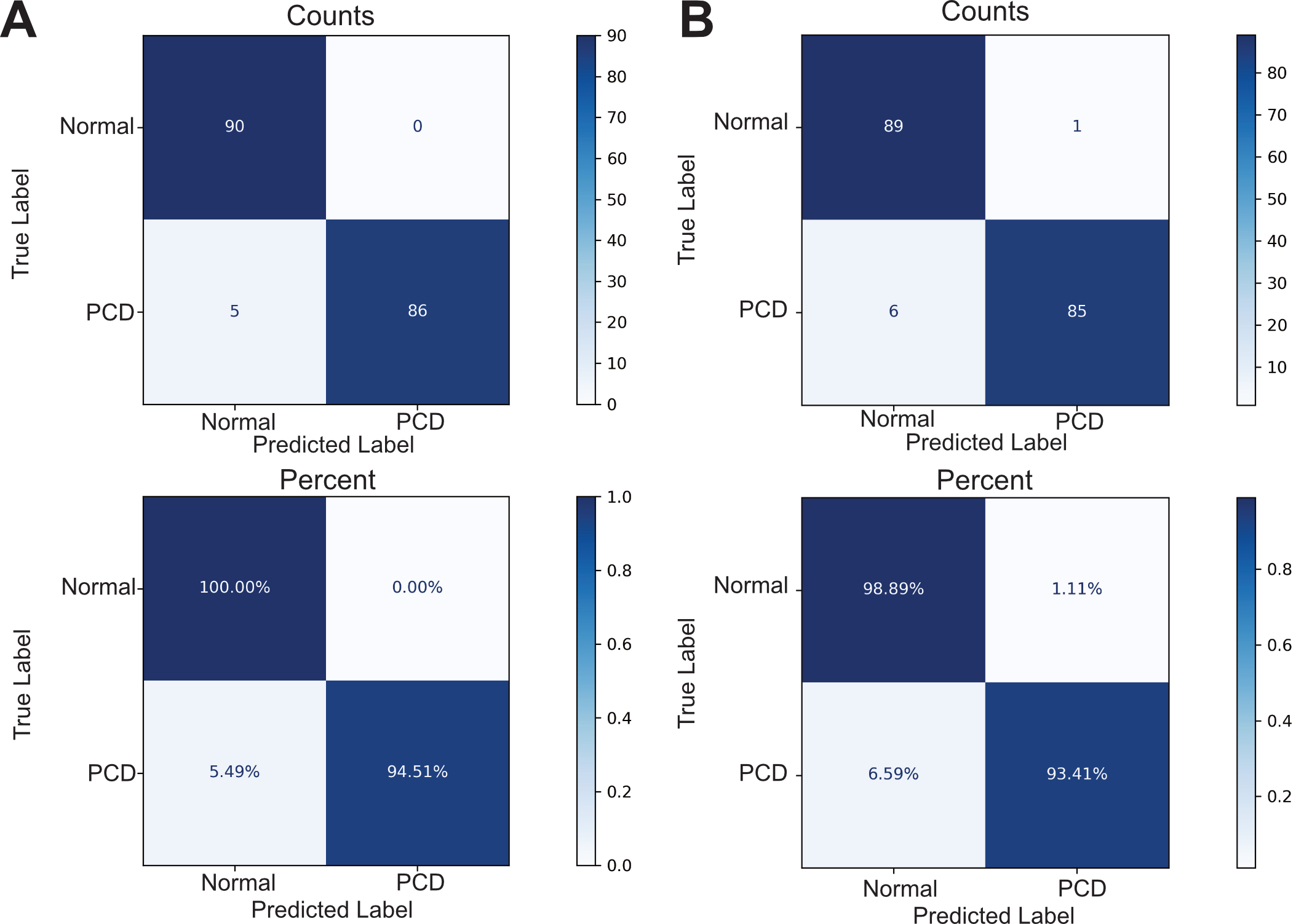

**(A)** Confusion matrices for the Random Forest classifier. Top panel shows the raw counts of prediction for control and PCD samples. The bottom panel shows the same result normalized by true condition. **(B)** Confusion matrices of the logistic regression classifier with top and bottom panel displaying the raw counts and percent normalized counts, respectively.

**Supplemental Figure S5. Misclassified CNN predictions.**

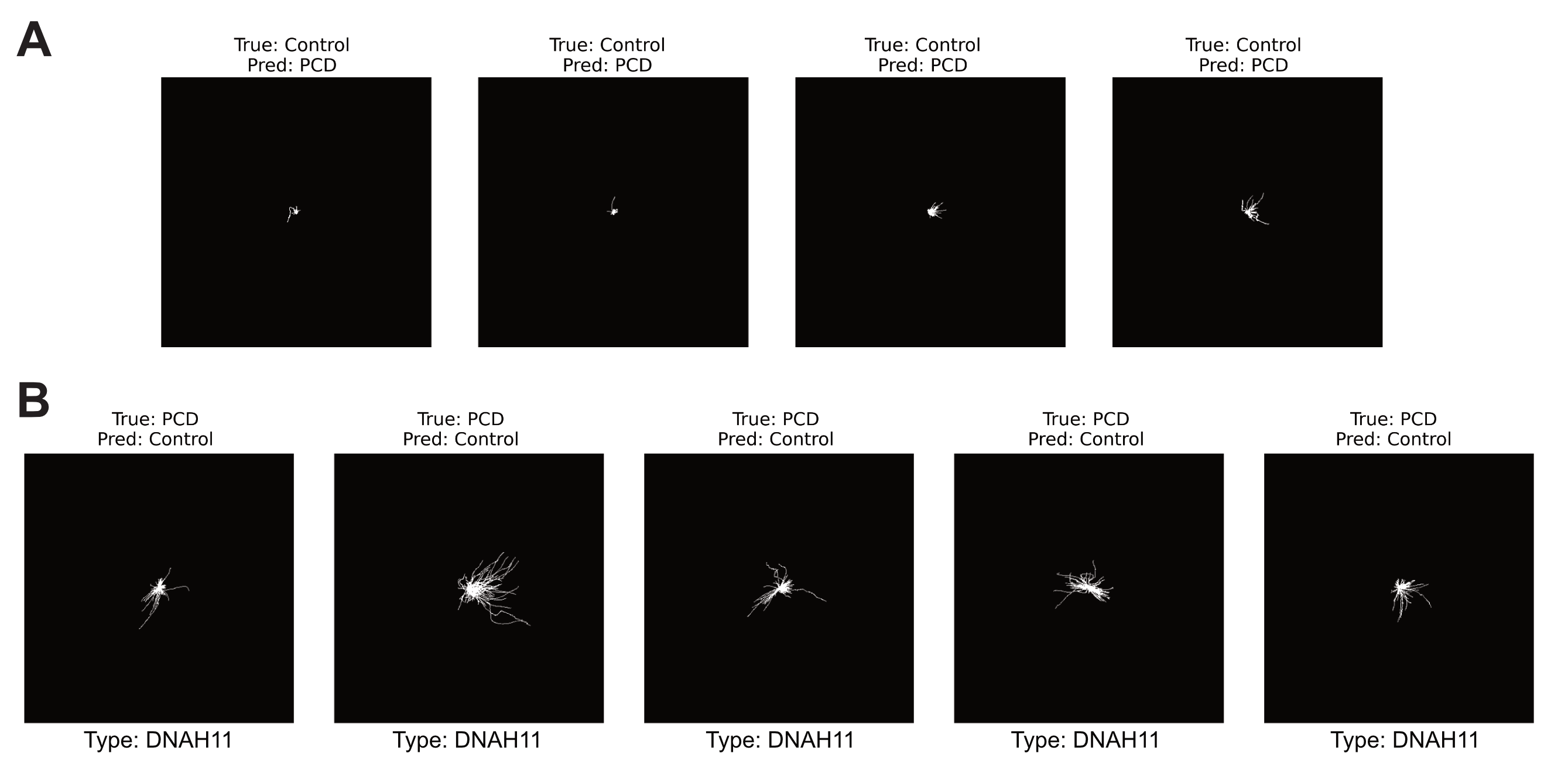

**(A)** Normal trajectory plot images that the CNN model misclassified as PCD (false positives). **(B)** False negative image classifications, where the model classified PCD plots as control.

**Supplemental Figure S6. Complete visual trajectory analysis of cystic fibrosis samples.**

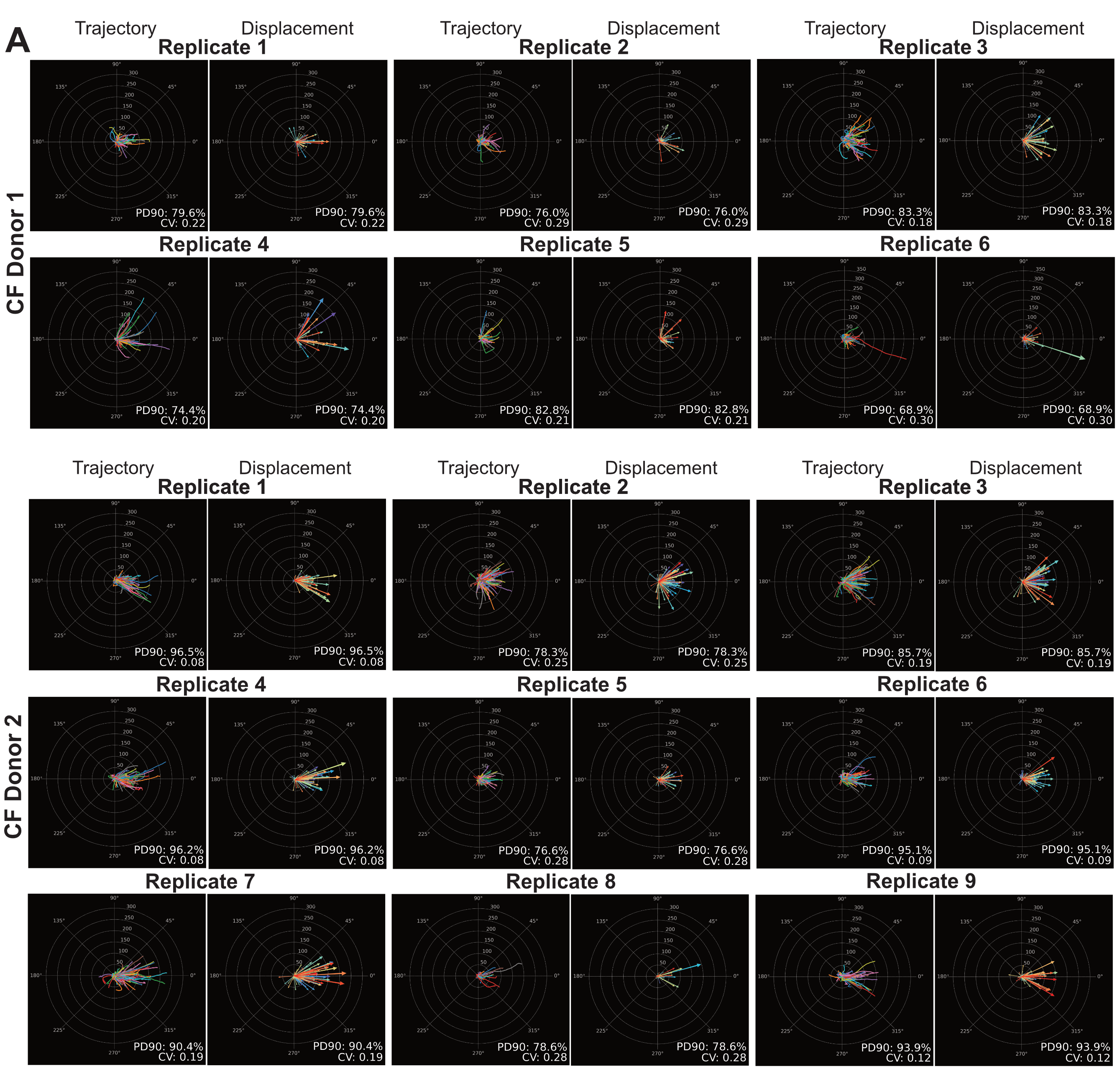

Trajectory and displacement plots displayed for all cystic fibrosis samples. Percent densest in 90 (PD90) and circular variance (CV) metrics are annotated for each plot.

**Supplemental Tables**

**Supplemental Table S1. Subjects’ genotypes.**

| **Donor group** | **Gene** | **Subject** | **Variant** |
| --- | --- | --- | --- |
| Normal |  | HTEC319 |  |
|  |  | HTEC323 |  |
|  |  | HTEC323 |  |
|  |  | HTEC328 |  |
|  |  | HTEC257 |  |
|  |  | HTEC298 |  |
|  |  | HTEC304 |  |
|  |  | HTEC316 |  |
|  |  | HTEC332 |  |
|  |  | HTEC325 |  |
| Heterozygous | *DNAH11* | WUPCD139 | cis c.4879C>T, c.7553T>C |
| Primary Ciliary Dyskinesia | *DNAAF5* | WU5012PCD1 | Homozygous c.2384T>C |
|  |  | WU5012PCD3 | Homozygous c.2384T>C |
|  |  | WU503PCD1 | Homozygous c.2384T>C |
|  |  | WU504PCD1 | Homozygous c.2384T>C |
|  |  | WUPCD501 | Homozygous c.2384T>C |
|  | *DNAH11* | WUPCD185 | c.518 A>G, c.8230C>T |
|  |  | WUPCD133 | c.6727C>T, c.13306G>A |
|  |  | WUPCD128 | c.3530T>C, c.5924+5G>A |
|  |  | INDPCD188 | c.921+3_921+6delAAGT, c.1934A>G |
|  | *RSPH1* | WUPCD169 | Homozygous c.85G>T |
|  | *DNAH5* | WUPCD186 | c.10815delT, c.C8404T |
|  | *DNAAF1* | WUPCD187 | c.707_708del, c.1415_1416del |
|  | *HYDIN* | WUPCD114 | c.10426T, c.4866delT |
|  | *RSPH4A* | INDPCD189 | c.6244C>T, c.13515_13526dup |
| Cystic Fibrosis | p.F508del | WUCF101 | Homozygous c.1521_1523de  l |
|  |  | WUCF102 | Homozygous c.1521_1523del |

**Supplemental Table S2. Statistical description of track metrics.**

|  | **Median** | | **Interquartile range** | |
| --- | --- | --- | --- | --- |
| **Feature** | **Normal** | **PCD** | **Normal** | **PCD** |
| Circular Variance | 0.30 | 0.64 | 0.14 – 0.57 | 0.48 – 0.81 |
| Confinement Ratio | 0.58 | 0.14 | 0.47 – 0.66 | 0.06 – 0.26 |
| Linearity of Forward Progression | 0.58 | 0.15 | 0.47 – 0.67 | 0.06 – 0.26 |
| Max Distance Traveled (μm) | 34.97 | 3.83 | 23.50 – 44.92 | 2.08 – 9.76 |
| Mean Directional Change Rate (radians/s) | 93.48 | 144.59 | 79.49 – 107.17 | 137.47 – 151.33 |
| Mean Straight Line Speed (μm/s) | 29.71 | 2.57 | 21.78 – 42.26 | 0.50 – 5.79 |
| Percent in Densest 90 (%) | 71.70 | 48.94 | 52.46 – 89.58 | 41.38 – 60.71 |
| Total Distance Traveled (μm) | 64.85 | 35.94 | 51.68 – 86.74 | 29.68 – 42.08 |
| Track Displacement (μm) | 34.05 | 3.39 | 22.61 – 44.30 | 1.57 – 9.24 |
| Track Max Speed (μm/s) | 271.00 | 83.97 | 187.53 – 413.84 | 28.38 – 196.13 |
| Track Mean Speed (μm/s) | 56.04 | 12.00 | 36.98 – 80.31 | 7.70 – 17.30 |
| Track Min Speed (μm/s) | 10.77 | 0.72 | 5.16 – 18.27 | 0.32 – 1.79 |

**Supplemental Table S3. Training and test set sample distribution.**

| **Analysis type** | **Condition** | **Training set** | **Test set** |
| --- | --- | --- | --- |
| Classical machine learning | Normal | 211 | 90 |
|  | PCD | 210 | 91 |
| CNN | Normal | 211 | 90 |
|  | PCD | 210 | 91 |

**Supplemental Table S4. Hyperparameter selection for the three models after 5-fold cross validation.**

| **Model** | **Hyperparameter** | **Best parameter** |
| --- | --- | --- |
| XGBoost | colsample_bytree | 0.6 |
|  | learning_rate | 0.05 |
|  | max_depth | 3 |
|  | n_estimators | 500 |
|  | subsample | 0.8 |
| Random Forest | max_depth | 5 |
|  | min_samples_leaf | 1 |
|  | n_estimators | 100 |
| Logistic Regression | C (inverse regularization strength) | 0.1 |
|  | penalty | l1 |

**Supplemental Table S5. CNN architecture in Pytorch.**

| **Layer group** | **Layer type** | **Parameters** | **Output shape** |
| --- | --- | --- | --- |
| Input | Input image | 1-channel grayscale | (1, 500, 500) |
| Convolutional block 1 | Conv2d | 16 filters, 3x3 kernel, stride 1, padding 1 | (16, 500, 500) |
|  | ReLU |  | (16, 500, 500) |
|  | MaxPool2d | 2x2 kernel, stride 2 | (16, 250, 250) |
| Convolutional block 2 | Conv2d | 32 filters, 3x3 kernel, stride 1, padding 1 | (32, 250, 250) |
|  | ReLU |  | (32, 250, 250) |
|  | MaxPool2d | 2x2 kernel, stride 2 | (32, 125, 125) |
| Convolutional block 3 | Conv2d | 64 filters, 2x2 kernel, stride 1, padding 1 | (64, 125, 125) |
|  | ReLU |  | (64, 125, 125) |
|  | MaxPool2d | 2x2 kernel, stride 2 | (64, 62, 62) |
| Classifier | AdaptiveAvgPool2d | Output size (1, 1) | (64, 1, 1) |
|  | Flatten |  | (64) |
|  | Linear (fully connected) | 128 output features | (128) |
|  | ReLU |  | (128) |
| Output | Linear (fully connected) | 2 output features (logits) | (2) |

**Supplemental Table S6. Training and test set grouped and separated by donors for model retraining.**

| Donor separated sets | Genetic variant or Normal | Number of samples | Number of donors |
| --- | --- | --- | --- |
| Training | Normal | 213 | 7 |
| (70%) | *DNAH11* | 69 | 3 |
|  | *DNAAF5* | 61 | 2 |
|  | *HYDIN* | 44 | 1 |
|  | *RSPH1* | 21 | 1 |
|  | *DNAAF1* | 12 | 1 |
| Test | Normal | 88 | 3 |
| (30%) | *DNAH11* | 45 | 1 |
|  | *DNAAF5* | 24 | 3 |
|  | *RSPH4A* | 13 | 1 |
|  | *DNAH5* | 12 | 1 |

**Supplemental Table S7. Model performance reevaluated on donor separated sets**.

| **Retrained Model** | **Class** | **Precision** | **Recall** | **F1-Score** | **Accuracy** |
| --- | --- | --- | --- | --- | --- |
| XGBoost | Normal | 0.98 | 0.97 | 0.97 | 0.97 |
|  | PCD | 0.93 | 0.98 | 0.97 |  |
| CNN | Normal | 0.95 | 0.93 | 0.94 | 0.95 |
|  | PCD | 0.94 | 0.96 | 0.95 |  |

**Supplemental Table S8. Genotypes and captured videos for CBF analysis.**

| **Sample group** | **Genetic variant** | **Videos analyzed** |
| --- | --- | --- |
| Unaffected Control | Not Applicable | 82 |
| Primary Ciliary Dyskinesia (PCD) | *DNAAF5* | 6 |
|  | *DNAH11* | 12 |
|  | *HYDIN* | 6 |
|  | *RSPH1* | 7 |
|  | *RSPH4A* | 3 |
|  | *DNAH5* | 24 |
|  | *DNAAF1* | 24 |
|  | Total samples | 164 |

**Supplemental Table S9. CBF logistic regression training and test set sample distribution**.

| Condition | Training set | Test Set |
| --- | --- | --- |
| Normal | 57 | 25 |
| PCD | 57 | 25 |

**Supplemental Table S10. False negative misclassifcations for XGBoost and CNN model.**

| **Model** | **Misclassification index** | **Genetic variant** |
| --- | --- | --- |
| XGBoost | 1 | *DNAH11* |
|  | 2 | *DNAH11* |
|  | 3 | *DNAH11* |
|  | 4 | *DNAH11* |
|  | 5 | *DNAH11* |
| CNN | 1 | *DNAH11* |
|  | 2 | *DNAH11* |
|  | 3 | *DNAH11* |
|  | 4 | *DNAH11* |
|  | 5 | *DNAH11* |
